# Functional traits and community composition: multilevel models outperform community-weighted means

**DOI:** 10.1101/183442

**Authors:** Jesse E. D. Miller, Ellen I. Damschen, Anthony R. Ives

**Affiliations:** Department of Zoology, University of Wisconsin, 430 Lincoln Dr., Madison, WI 53706 USA; Current address: Department of Environmental Science and Policy, University of California, Davis, 1 Shields Avenue, Davis, California 95616 USA

**Keywords:** Community assembly, ecological community, environmental gradients, linear mixed models, Whittaker Siskiyou Mountains data

## Abstract

1. Plant functional traits are increasingly being used to infer mechanisms about community assembly and predict global change impacts. Of the several approaches that are used to analyze trait-environment relationships, one of the most popular is community-weighted means (CWM), in which species trait values are averaged at the site level. Other approaches that do not require averaging are being developed, including multilevel models (MLM, also called generalized linear mixed models). However, relative strengths and weaknesses of these methods have not been extensively compared.
2. We investigated three statistical models for trait-environment associations: CWM, a MLM in which traits were not included as fixed effects (MLM1), and a MLM with traits as fixed effects (MLM2). We analyzed a real plant community dataset to investigate associations between two traits and one environmental variable. We then analyzed permutations of the dataset to investigate sources of type I errors, and performed a simulation study to compare the statistical power of the methods.
3. In the analysis of real data, CWM gave highly significant associations for both traits, while MLM1 and MLM2 did not. Using *P*-values derived by simulating the data using the fitted MLM2, none of the models gave significant associations, showing that CWM had inflated type I errors (false positives). In the permutation tests, MLM2 performed the best of the three approaches. MLM2 still had inflated type I error rates in some situations, but this could be corrected using bootstrapping. The simulation study showed that MLM2 always had as good or better power than CWM. These simulations also confirmed the causes of type I errors from the permutation study.
4. The MLM that includes main effects of traits (MLM2) is the best method for identifying trait-environmental association in community assembly, with better type I error control and greater power. Analyses that regress CWMs on continuous environmental variables are not reliable because they are likely to produce type I errors.

## Introduction

Analyzing plant functional traits is an increasingly popular approach for understanding plant community assembly (Cornwell & Ackerly 2015; Funk *et al.* 2017). Functional traits are relatively easy-to-measure characteristics of plants representing key life history processes that are difficult to measure directly (Lavorel & Garnier 2002; Reich 2014). Functional traits may facilitate understanding mechanisms by which community composition responds to environmental gradients and thus provide inference beyond species-focused investigations of community assembly processes (Lavorel & Garnier 2002; Funk *et al.* 2017). However, testing how community-level functional traits vary in response to ecological gradients poses an analytical challenge, since environmental gradients are measured at the plot or site level (hereafter site) and functional traits are often measured at the species level. This means that somehow, values of functional traits for many species must be integrated and compared to single environmental variable values. Several statistical approaches to this challenge have been developed, but there has been little comparison of different methods and their relative strengths and weaknesses. To ensure their reliability, methods should be compared for standard statistical properties such as type I error control and statistical power.

One of the most common approaches for analyzing trait-environment relationships is to use a “community-weighted mean” (CWM; Ricotta and Moretti 2011), which is the average of all trait values at each site, weighted by species abundance. A recent “trait-based ecology” review highlights the common usage of the CWM (Funk *et al.* 2017). Indeed, we found 188 papers in the peer-reviewed literature (Web of Knowledge, 7 March 2017, papers that contain the term “community-weighted mean*” as topics). One appeal of the CWM approach is that relationships between CWM values and environmental gradients can be analyzed using simple regression. However, CWM reduces a large amount of data to a single trait value at each site, which raises concerns about possible loss of statistical power, since statistical power is related to sample size. Furthermore, because multiple sites contain the same species, CWM values for different sites along the gradient are not independent, and the loss of species-level information makes it impossible to account for this non-independence in statistical analyses. Typical CWM analyses ignore this non-independence, which is likely to cause problems with type I error control (Ives & Zhu 2011).

An alternative approach to CWMs is multilevel models (MLM, also known as linear mixed models; Pollock *et al*. 2012; Jamil *et al*. 2013). In this approach, the occurrence (1 or 0) or abundance of each of *n* species in each of *m* sites is used as the response variable. Each species × site combination is assigned a species-level trait and a site-level environmental variable. The test for functional traits being associated with environmental gradients involves the trait-by-environment interaction. Because the *n* × *m* species-site data points are not independent (because species occur at multiple sites), random effects for species are included to allow species to have unique responses to the environmental gradients. Thus, unlike CWM, MLMs explicitly account for non-independence of data among sites. Nonetheless, MLMs can be constructed to include different terms as predictor variables, and the choice of models can affect the statistical conclusions about the significance of the trait-by-environment interactions.

In this study, we compare the performance of three approaches to analyzing trait-environment relationships. We first analyze real plant community data using CWM and two MLMs: MLM1 that was proposed by Pollock *et al*. (2012), and MLM2 that includes a fixed effect for traits (Jamil *et al*. 2013). We then use permutation and simulation studies to investigate any variation in results among methods, test for sources of possible type I errors, and compare the statistical power. Our goal is to assess which methods provide statistically robust tests of trait-environment associations.

## Materials and Methods

### Dataset

To test the ability of the different approaches to determine associations between traits and the environment, we used a community dataset from a subset of Robert Whittaker’s historic plant community study sites in the Siskiyou Mountains of Southwest Oregon (Whittaker 1960; Damschen *et al*. 2010). Within this dataset, we selected low-elevation, non-serpentine sites for study. Vegetation surrounding the sites consists primarily of conifer forest dominated by Douglas fir (*Pseudotsuga menziesii*), with an assortment of conifer and evergreen hardwood trees and a diverse herbaceous understory. Community surveys were conducted in the summer of 2007 following Whittaker’s original sampling methods (Whittaker 1960; Damschen *et al*. 2010). Whittaker chose sites to represent the full range of topography, with sites distributed across each of the ten topographic moisture levels. A single 0.1 ha study plot was established at each study site, and 25 1 x 1-m quadrats were established along a 50-m transect running down the center of each plot. Species abundances were calculated from the number of 100 quadrat corners each species intercepted. For our analyses, we limited the data to species that intercepted at least one quadrat corner so that all plants had an abundance ranking.

We chose two functional traits, leaf carbon-to-nitrogen ratio (C:N) and plant vegetative height (height), and one environmental variable, Whittaker’s topographic moisture gradient (TMG). The two chosen functional traits relate to plant resource acquisition life history strategies (Cornelissen *et al*. 2003; Poorter & Bongers 2006). The C:N ratio is often considered a surrogate of competitive ability; plants with lower C:N may grow faster, but may have lower stress tolerance (Poorter & Bongers 2006). Similarly, taller plants can shade out neighbors, but shorter plants may be more stress tolerant (Kunstler *et al*. 2015). The topographic moisture gradient ranks each site on a scale from 1-10, where sites on more mesic, north-facing slopes receive lower numbers, and sites on warmer, south-facing slopes receive higher numbers. We used a standardized version of the topographic moisture gradient following Damschen *et al*. (2010). Ecologically, we expected that leaf C:N ratios and height would both decline with an increase in the TMG score, that is, as sites become drier and warmer.

### The community-weighted-mean (CWM) model

We calculated CWM for each of the two traits at each site by multiplying trait values for the species that occur at the site by their abundances, adding these values, and then dividing by the sum of species abundance values. Community-weighted means were then regressed against TMG scores.

### Multi-level model without fixed traits (MLM1)

MLM1 is the multilevel model proposed by Pollock *et al*. (2012). MLMs use long-form community data, with a separate row *i* for each potential species-site combination. We formulated MLM1 for binomial data, with the abundance of each species taking a value from 0 to 100 in each site. The MLM is thus a logit-normal binomial generalized linear mixed model (GLMM) with the form

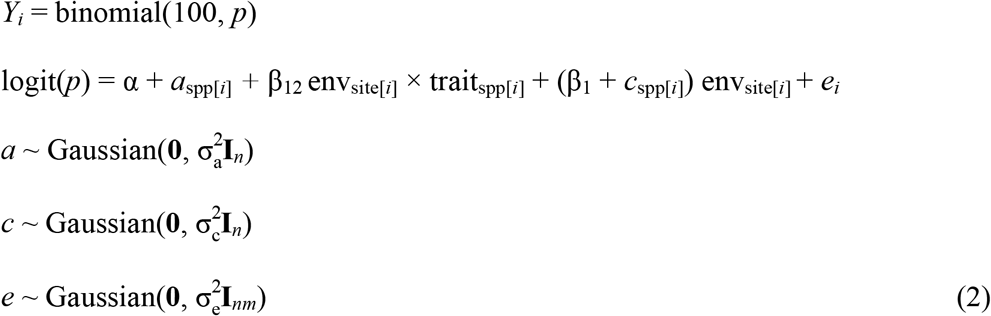

Here, *Y_i_* (*Y_i_* = 0,…, 100) is the observed abundance for each of the *i* = *n* × *m* species-site data rows. Following the convention of multilevel models (Gelman & Hill, 2007), the functions spp[*i*] and site[*i*] map row *i* onto the corresponding species and sites, so that trait_spp[*i*]_ gives the trait (C:N or Height) of the species in row *i* and env_site[*i*]_ gives the topographic moisture index. The fixed effect α gives the overall average abundance of species among sites, and the fixed effect β_1_ gives the mean response to the environmental gradient env. Random effect *a*_spp[*i*]_ allows different species to have different overall abundance, and random effect *c*_spp[*i*]_ allows different species to have different responses to TMG (environmental gradient). Random effect *e* gives observation-level variance which accounts for greater-than-binomial variance in the observations. The association between traits and the environmental gradient (topographic moisture gradient here) is given by the fixed effect β_12_, and this is the target of statistical testing.

### Multi-level model with fixed traits (MLM2)

MLM2 is similar to MLM1 but follows Jamil et al. (2013) by including an additional fixed effect β_2_ for traits on mean species abundance and an additional random effect for site:

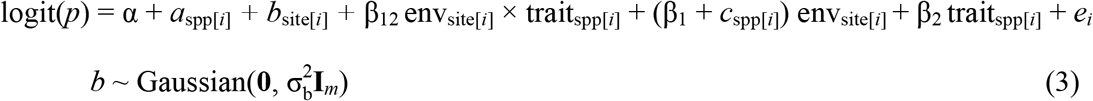

These additional terms in MLM2 force less of the variation in species abundance onto the trait-by-environment interaction term β_12_ compared to MLM1. Specifically, the additional fixed effect β_2_ allows traits to affect species abundances without doing so through the interaction with the specific environmental gradient considered in the equation. The random effect *b* allows sites to differ in mean species abundance independently from the environmental gradient.

Both MLM1 and MLM2 can be performed as either univariate models (one trait and one environmental variable in a model) or multivariate models (one or more traits or environmental variables). For simplicity, we present univariate analyses for single traits with our single environmental variable in the main text and analyses of multivariate models with both traits and the environmental variable in the Appendix.

### Permutations

We performed three permutation analyses designed to expose different properties of the data and statistical methods:

1. permuting trait values among species while maintaining the distribution of species among sites (Fig. 1B)
2. permuting values of the environmental variable among sites while maintaining within-site species composition (Fig. 1C)
3. permuting species composition along with their traits among sites while maintaining site-level species richness (Fig 1D).

**Figure 1.**
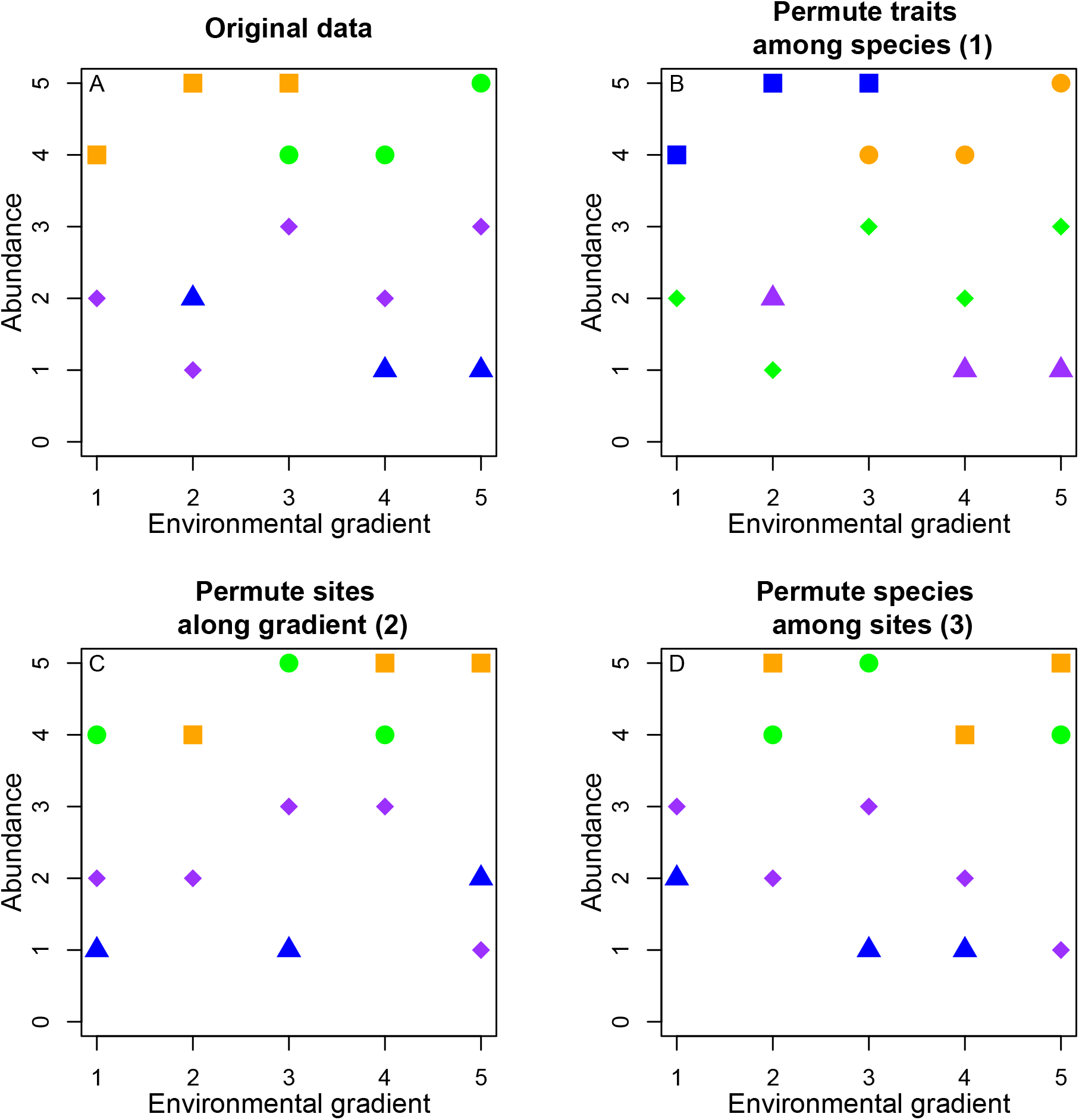
Conceptual illustration of plant communities along an environmental gradient and examples of the three permutations. Each point represents a species at a site; shapes represent species identity and colors represent trait values. Panels show (a) original data before permutation, (b) permuting traits among species, (c) permuting the environmental variable among sites, and (d) permuting species among sites.

Each permutation randomizes the association between traits and environmental gradient, but they differ what characteristics of the communities they preserve. For example, the first permutation breaks down the association between traits and environmental gradient while preserving any association between the environment and species richness, and also any non-random association of species with the environment. The ideal statistical test, when applied to data generated by all three of these permutations, would show correct type I errors, rejecting the null hypothesis of no association at the nominal alpha significance level. If one or more of these permutations generates inflated type I errors, we can compare them to identify the source of inflated type I error.

Of the three permutations, the first permutation corresponds to the null hypothesis that is generally being tested in studies using CWM: the functional traits do not explain the relative abundances of species among communities. The second permutation preserves the structure of the communities, changing only the environmental gradient among communities. The third permutation preserves the number of species per site and also maintains as much as possible the same mean occurrence of each species among sites. For each site, species were selected at random in proportion to their occurrences among all sites. Thus, if a species occurred in 4 sites, it was selected with twice the probability of a species occurring in 2 sites. For each analysis, we performed 1000 permutations and tabulated the proportion of permutation datasets for which the null hypothesis β_12_ = 0 was rejected at the alpha significance level of 0.05. If the statistical tests have proper type I error control, the null hypothesis should be rejected in 1/20 of the datasets.

### Simulations

We performed a simulation study to test the statistical power of the different analysis approaches. We built our simulation study around the fit of MLM2 to the Whittaker dataset because MLM2 gave a better fit than MLM1 (likelihood ratio test, logLik(MLM2) = –3163.8, logLik(MLM1) = –3196.8, χ^2^2 = 66, *P* << 0.0001). We considered three simulation scenarios for equation (3): (i) β_2_ = 0, (ii) σ_a_ = σ_c_ = 0, and (iii) β_2_ = σ_a_ = σ_c_ = 0. These correspond roughly to permutations (1) through (3). The three simulation scenarios impose restrictions in addition to the null hypothesis of the statistical tests that β_12_ = 0. In simulation (i) there is no effect of trait values on the mean abundance of species among sites, yet there is variation among species in their response to the environmental gradient. Like permutation (1), simulation (i) assumes that trait values do not determine the abundance of species; nonetheless, the environment can play a role in the variation in abundance of species among sites that does not involve the trait. In simulation (ii), species do not differ in either mean abundance or their trait-independent response to the environmental gradient. Like permutation (2), the environmental value at a site does not determine the abundance of species at that site, although there could be other site-specific factors that generate variation in the abundances of species among sites. Simulation (iii) combines the restrictions of simulations (i) and (ii). Like permutation (3), in simulation (iii) neither environment nor traits explain the abundance of species among sites.

For each simulation scenario, we fit MLM2 to the dataset under the assumptions (i)-(iii) and then simulated 1000 datasets from the fitted model. Occasionally datasets were produced where one or more species did not occur in any sites; when this occurred, we set an abundance of 1 in the site that had the highest abundance of that species in the real dataset.

For each simulation dataset, we applied the three models and tabulated the proportion of simulated datasets for which the null hypothesis β_12_ = 0 was rejected at the alpha-significance level of 0.05. In the simulations when β_12_ is set to 0, the analysis gives type I error rates. To investigate the statistical power of the methods, we set β_12_ = 0, 0.1, 0.2, 0.3, 0.4, and 0.5 in the simulations and tabulated the proportion of datasets for which β_12_ = 0 was rejected; provided type I error control is adequate (i.e., 5% of the simulated datasets are rejected when β_12_ = 0), greater rejection rates as β_12_ increases implies greater power.

Finally, we wanted to determine whether the inclusion of the fixed effect for traits in MLM2 (β_2_ in equation 3) was responsible for performance differences between MLM1 and MLM2, rather than the random effect for site (σ_b_ in equation 3). Therefore, we performed the simulations with MLM2 setting σ_b_ = 0, and we also set σ_b_ = 0 when fitting MLM2. This means that in the simulation study, MLM1 and MLM2 only differ by the inclusion of β_2_ in MLM2.

All analyses were performed in R version 3.3.2 (R Core Team 2016). The R code for MLM2, and the analyzed dataset, are available in the Appendix.

## Results

### Observed data

In tests of association between TMG and both functional traits (C:N and Height), the relationships between the environmental variable and traits in the CWM model were highly significant, while the MLM1 and MLM2 tests were not (Table 1, S1-S3). Multivariate formulations of MLM1 and MLM2 were qualitatively similar to univariate formulations (Table S4). This contrast between CWM and multilevel models begs the question of which is correct, so we simulated data with the fitted MLM2 (including the estimate of the trait-by-environment interaction β_12_) 1000 times and refit each model. The distribution of the resulting 1000 trait-by-environment coefficients (slopes for CWM and β_12_ for MLM1 and MLM2) was then used to calculate confidence intervals. We simulated data using MLM2 because this model fit the data better than MLM1 (likelihood ratio test, χ^2^2 = 66, *P* << 0.0001). Note that for MLM2, this simulation procedure is a parametric bootstrap for tests of the model coefficients. For MLM1 and MLM2 the simulated *P*-values were close to the *P*-values calculated from the GLMMs, but for CWM the simulated *P*-values were much higher and non-significant. This implies that CWM results for the real dataset were false positives and that CWM has very poor type I error control.

**Table 1.** Model *P*-values for tests of whether two functional traits predict species abundances along TMG under three methods: CWM, MLM1, and MLM2. To assess the validity of the *P*-values obtained from the three methods, we also performed simulations using the fitted MLM2. The *P*-values from simulations are the proportion of 1000 simulations for which the trait-by-environment interaction term (slope for CWM and β_12_ for MLM1 and MLM2) was less than zero (multiplied by 2 to give a 2-tailed test). The simulated *P*-values are correct, in the sense that they properly account for the non-random structure of the data given by MLM2.

|  | CWM |  | MLM 1 |  | MLM 2 |  |
| --- | --- | --- | --- | --- | --- | --- |
| Trait | C:N | Height | C:N | Height | C:N | Height |
| $P$-value | **0.0018** | **<0.001** | 0.674 | 0.613 | 0.095 | 0.06 |
| Simulation $P$-value | 0.402 | 0.268 | 0.257 | 0.210 | 0.105 | 0.062 |

### Permutations

We performed three permutations to identify the source of poor type I error control in the CWM model and also evaluate the performance of MLM1 and MLM2. Permutation (1) of traits among species yielded highly inflated type I error rates for the CWM model: the null hypothesis was rejected 38% and 39% of the time at the alpha significance level of 0.05 for both traits (C:N and Height). In contrast, both the MLM1 and MLM2 models rejected considerably less than 5% of the permuted datasets. For permutation (2) of the environmental variable among sites, CWM regression yielded good type I error control. However, the MLM1 and to a lesser extent MLM2 models had inflated type I error rates for C:N. For permutation (3) of species (and their traits) among sites, the CWM model had good type I error control, while both the MLM1 and MLM2 models rejected the null hypothesis considerably less frequently than they should. Multivariate tests with the MLM1 and MLM2 models that contained both traits yielded qualitatively similar results to the univariate formulations (Table S5). In summary, the CWM model had very poor type I error control under permutation (1), MLM1 and to a lesser extent MLM2 models had poor type I error control under permutation (2), and all models had good type I error control under permutation (3).

Permutation (1) of traits among species should yield random data with respect to the null hypothesis of no trait-by-environment association, yet CWM rejects this null hypothesis for the permutation datasets far too often. The poor type I error control of CWM is caused by the lack of independence of CWM values among sites that contain the same species. Some species occur in many sites (Fig. 2), and therefore the sites where they occur are more likely to have the same CWM values. This is exacerbated by a few species having very high mean abundances, and these species will contribute more heavily to the CWM values (Fig. 2). From the permutation datasets, we calculated the correlation among CWM values between sites. For permutation (1) of traits among species, the correlations of CWM were predominantly positive and increased for pairs of sites that shared a greater proportion of their species, indicating that the non-independence is driven by sites sharing the same species. In contrast, for permutation (2) of environments among sites and permutation (3) of species among sites, the correlations were on average zero and were never strongly positive or negative. This independence of CWM values under permutations (2) and (3) explains the correct type I error control of CWM under these permutations.

**Figure 2.**
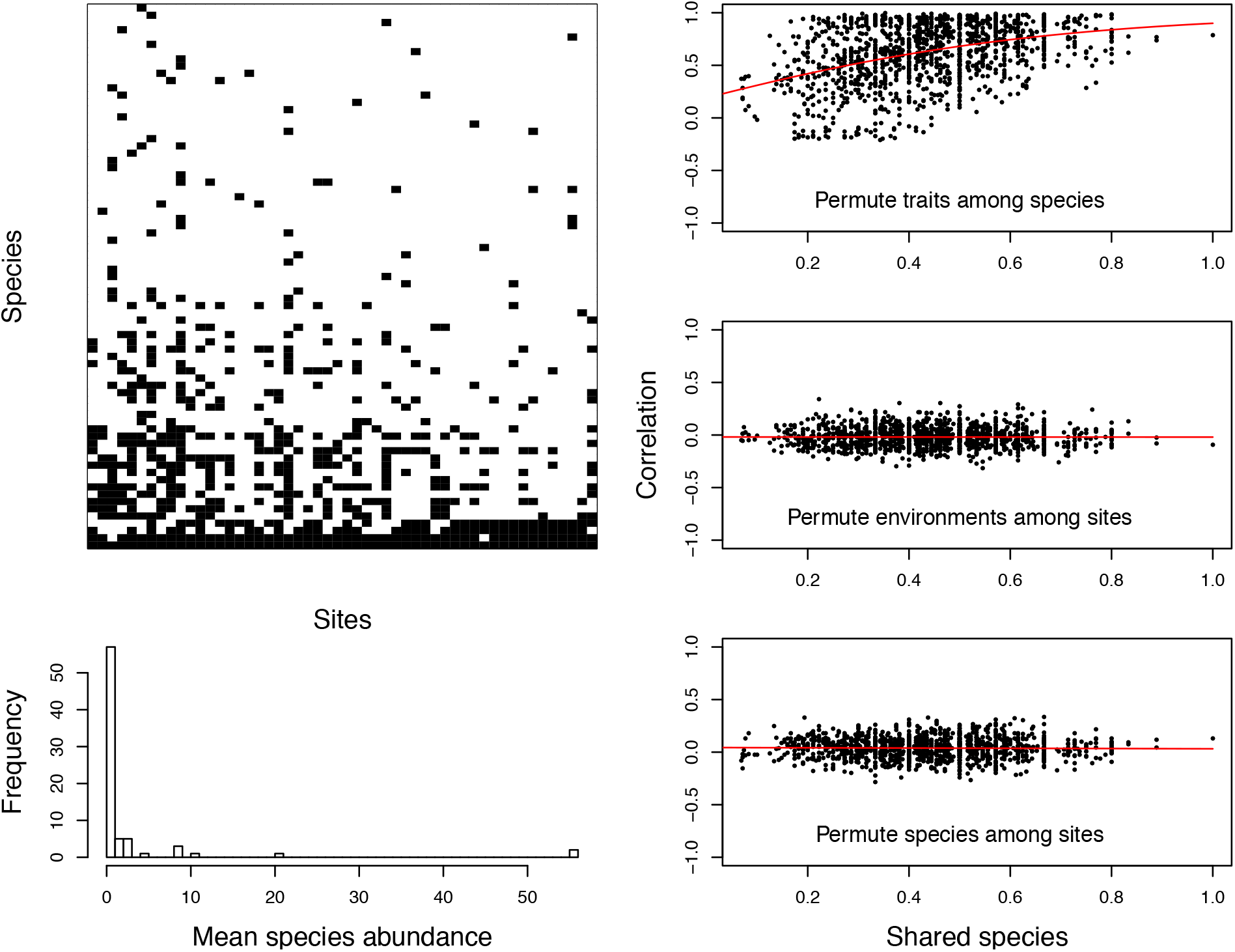
Identifying the cause for the inflated type I error rates of CWM regression. The top left panel gives the presence of species among sites (black), with sites sorted by increasing TMG (environmental gradient) and species sorted by the number of sites in which they occur. The lower left panel gives the frequency distribution of mean species abundances. The panels on the right give, for each of the three permutations, the relationship between the pairwise correlation between CWM values for each site and the proportion of shared species between the sites. The correlation in CWM values was calculated from 100 permutations, and the proportion of shared species was 2 *(number of species shared by sites *i* and *j*)/((number of species in site *i*) + (number of species in site *j*)). The red lines give the function 2/(1 + exp(a + b*x)) fitted with least squares. For the permutation of environments among sites, site identity was based on the environmental values in sites, so the permutation moved species and their traits among sites. The high correlations of CWMs when traits are permuted among species (top-right panel) show the non-independence of CWM values among sites; this non-independence is worse between sites that share more species.

### Simulations

To test the statistical power of the different modeling approaches, we designed the simulation study to correspond to the permutation study. In simulation (i), trait values of a species did not affect its mean abundance among sites (Eq. 3, β_2_ = 0), yet species differed in overall abundance (σ_a_ > 0) and in their response to the environment in sites (σ_c_ > 0). CWM had very high type I rejection rates, while MLM1 and MLM2 had acceptable if low type I error rates (Fig. 3). These results are consistent with permutation (1). In this simulation, variation among species in either overall abundance or their response to the environment will cause the CWM values among sites to be non-independent. For example, if there is a species that has high abundance at high TMG values, then its trait value will weight heavily in CWMs for sites at the high end of TMG. If another species has high abundance at low TMG values, its trait value will weight heavily at the other end of TMG. This non-independence will then increase the chances of a large slope of CWM against TMG. Because the simple regression of CWM against the environmental gradient does not account for this non-independence, the test statistic assumes less variation in the slope and hence is more likely to incorrectly reject the null hypothesis of slope = 0.

**Figure 3.**
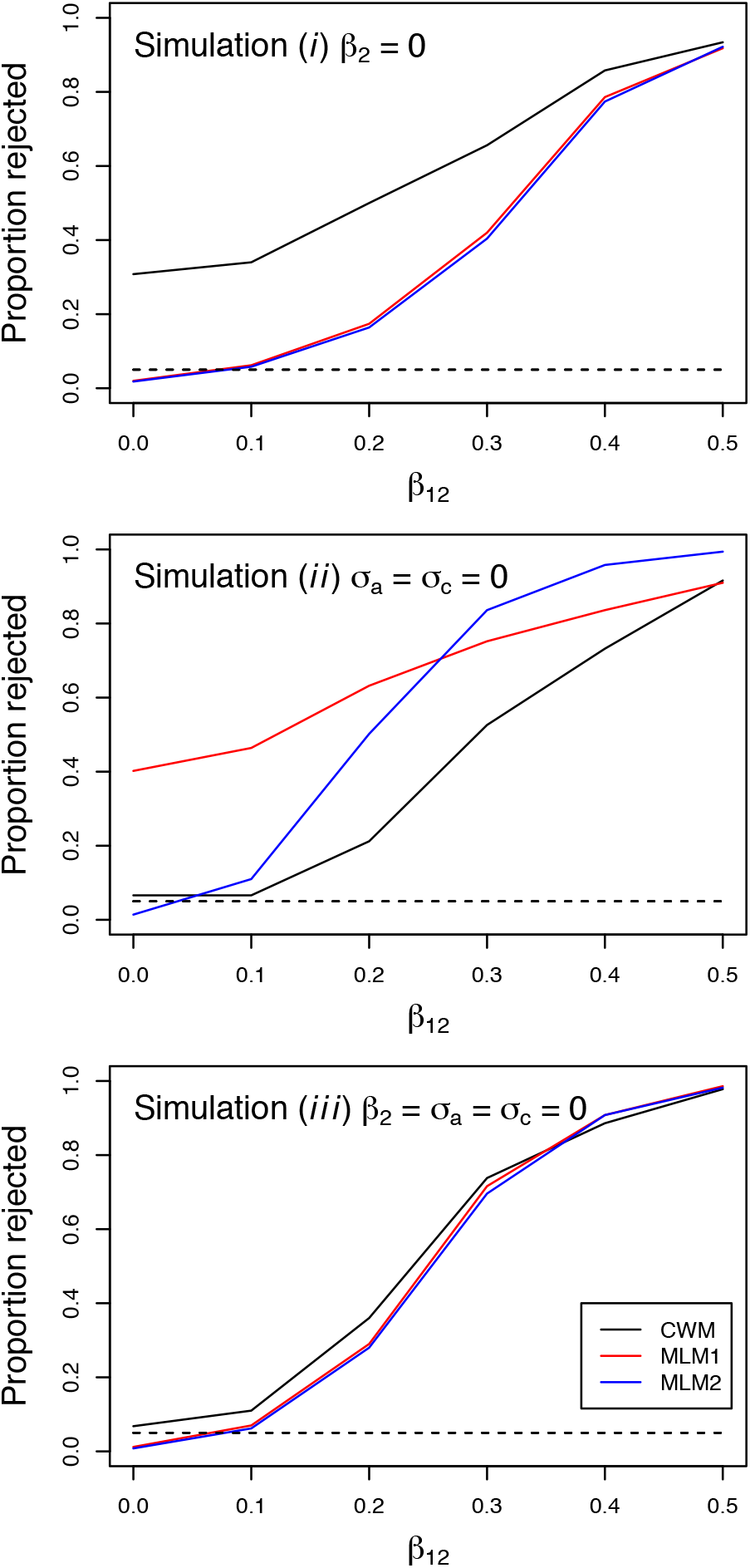
Results from simulation studies using equation (3) under scenarios: (i) β_2_ = 0, (ii) σ_a_ = σ_c_ = 0, and (iii) β_2_ = σ_a_ = σ_c_ = 0. MLM2 was first fit to the dataset under the three scenarios with β_12_ = 0. The fitted MLM2 was then used to simulate 1000 datasets at β_12_ = 0, 0.1, 0.2, 0.3, 0.4, and 0.5. CWM regression (black), MLM1 (red), and MLM2 (blue) were fit to each simulated dataset, and the proportion of the 1000 datasets for which the null hypothesis of no trait-by-environment interaction was rejected is plotted for each scenario.

In simulation (ii), there is no random variation in species mean abundance (σ_a_ = 0) and species do not differ in their responses to the environment (σ_c_ = 0), yet species trait values affect mean species abundances (β_2_ > 0). In this case, CWM values among sites are close to independent, because even though they might contain the same species, all species are similar to each other. However, MLM1 has poor type I error control. MLM1 does not include a term for a possible effect of traits on the mean abundance of species. Therefore, if there is an effect of traits on mean abundances, then the resulting variance is forced on the environment-by-trait interaction (Eq. 2, β_12_) because this is the only term in the model that contains a trait effect. In principle, this variation could be absorbed into the random effect *a* for differences among species in mean abundance that do not depend on trait values. However, in practice this did not happen for simulations based on this dataset. In contrast to the MLM1 model, the MLM2 model had type I error rates below the alpha significance level of 0.05. Despite this, the type I error rate increased rapidly with increases in β_12_, showing that MLM2 had considerably higher statistical power than the CWM model.

In simulation (iii), species have the same mean abundances (β_2_ = 0, σ_a_ = 0) and the same response to the environment (σ_c_ = 0). In this case, all three methods performed similarly. The MLM1 and MLM2 models had low type I error rates and, as a consequence, had slightly lower statistical power than the CWM model.

## Discussion

### Comparison of statistical methods

The three different statistical methods for analyzing functional trait responses to environmental gradients—community-weighted means and two multilevel models—vary substantially in performance. When analyzing the same dataset, CWM regression indicated highly significant trait-environment associations while MLM1 (Pollock *et al*. 2012) and MLM2 (Jamil *et al*. 2013) indicated no relationship between the same variables. This reflected the inflated type I error rates (false positives) that CWM regressions can produce because CWM values are not independent among sites. This lack of independence in CWM values is caused by species occurring in more than one site. If, for example, a species is common (and therefore weights heavily in the CWM) at several sites that occur at one end of the environmental gradient, then whatever trait value this species has will have high influence on the slope of the CWM regression. Because the CWM regression does not take into account the lack of independence among CWM values, it will ascribe greater significance to the regression slope than appropriate.

The MLM that does not contain traits as fixed (or random) effects (MLM1) also exhibits inflated type I error rates. This occurs when species mean abundances across all sites are not independent of their trait values (permutation 2, simulation ii). Because MLM1 does not account for this variation in overall mean abundances, it attributes the variation to the only term in the model that contains trait values, the trait-by-environment interaction term. If species are distributed among sites independently of the environmental gradient and have abundances that are independent of their trait values (permutation 3, simulation iii)—a situation that would be unlikely in real datasets—then all three methods show appropriate type I error rates. Because MLM2 is as good as or substantially better than the other approaches in all permutation and situation studies, we recommend it for general use in studies of trait-environment interactions.

The simulation study makes it possible to compare the power of the methods to detect trait-by-environment interactions when they have acceptable type I error control (Fig. 3). In simulation (iii), all three methods had similar power. In simulation (i) in which traits did not affect the overall abundance of species, MLM1 and MLM2 had similar power (while CWM had unacceptable type I error control). In simulation (ii) in which species did not have species-specific variation in responses to the environmental gradient, MLM2 had much higher power than CWM regression. This low power of CWM regressions might occur because CWM weights species according to their abundances, while MLM2 weights species equally; therefore, MLM2 will use the full variation among species in their responses to the environmental gradient.

### Calculating MLM P-values using bootstrapping

MLM1 had inflated type I error rates when the overall abundance of species across sites varied with trait values (simulation ii). MLM2 included a term for trait values affecting overall species abundances, and it consequently had good type I error control in simulation (ii). MLM2 still had slightly inflated type I error rates in permutation (2) for one trait (C:N in Table 2). Also, in permutations and simulations both MLM1 and MLM2 often had rejection rates that were below the 5% expected with a alpha confidence level of 0.05; these low rejection rates will decrease the power of the tests, but they will not lead to false claims of significance. We suspect that the source of low rejection rates is the nonlinearity of the link function used for the GLMMs and the difficulty this poses for estimating an interaction term. In linear mixed models (LMMs), if the interaction term is zero, the slopes corresponding to two predictor variables do not change; therefore, a non-zero interaction term is the only source of changes in slopes. In the logit-normal binomial MLM1 and MLM2, slopes change at different values of an independent variable (environment or trait values) even if there is no interaction term. Therefore, the statistical challenge is not simply showing that a slope changes, but instead that the slope changes in the way implied by the logit link function. In simulations using the MLM2 equation (3) with Gaussian data and a linear link function, the rejection rates were correct (results not presented).

**Table 2.**
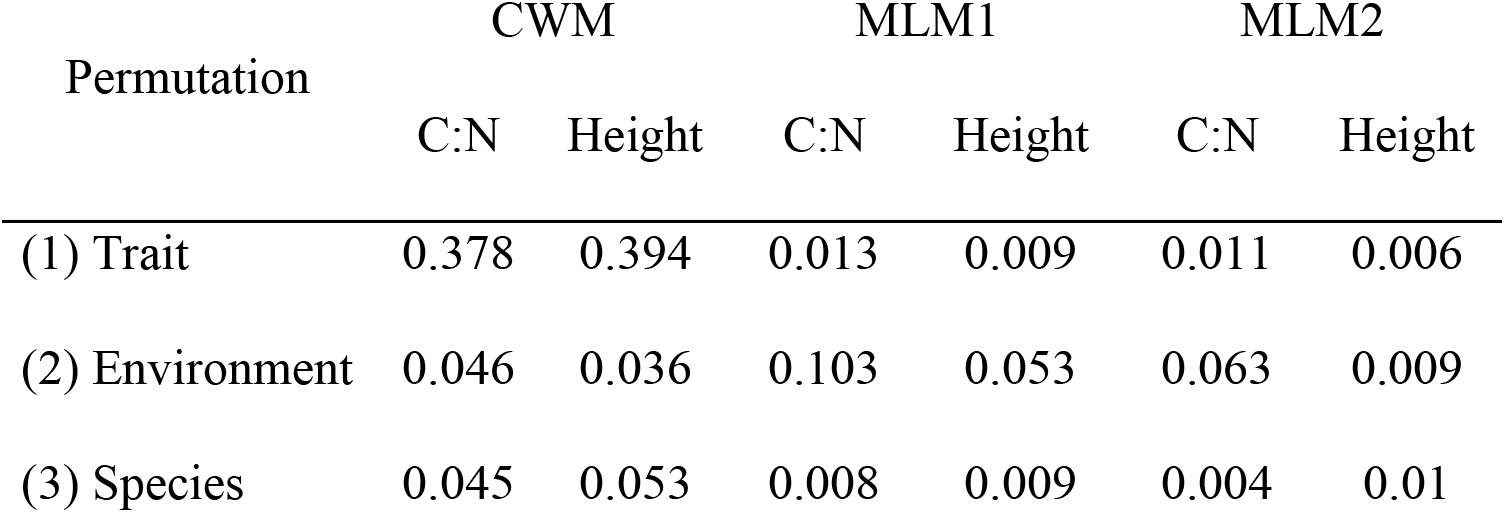
Proportion of permutation datasets for which each of the three models rejected the null hypothesis at a nominal alpha level of 0.05. Interactions between two traits (C:N and Height) and the environmental variable TMG were tested with three methods: CWM, MLM1, and MLM2.

To give correct *P*-values for GLMMs, we suggest using a bootstrap in which the fitted model is used to simulate a large number (e.g., 2000) of bootstrap datasets, and then each bootstrap dataset is then refitted by the same model. The resulting large number of estimates of the trait-by-environment interaction term β_12_ give the approximate distribution of the estimate of β_12_ which can be used for confidence intervals (e.g., the 95% bounds of the bootstrapped estimates of β_12_) and hypothesis tests (e.g., the proportion of estimates of β_12_ less than zero). These bootstrap tests of trait-by-environment interactions will give correct type I error rates and power.

### Suggestions for practitioners

CWM regression is the most commonly used method for studying community-level trait responses to environmental gradients. While CWM accurately represents the direction of trait-environment relationships (e.g., increasing plant height in response to increasing site moisture), interpreting *P*-values is problematic. We show how CWM can produce *P*-values < 0.05 nearly 40% of the time when trait values are randomly assigned to species. This problem is exacerbated when species differ in average abundance across sites, and if species richness decreases along the gradient—both common situations in community datasets. The degree to which CWM regressions are flawed depends on the properties of the specific dataset, and therefore it is difficult to know how badly inflated are the type I errors. Better performance might be expected when beta diversity is high, and when communities are not dominated by a few common species, because such situations will decrease the correlations of CWM values among sites. However, this is no guarantee. Many of the studies that use CWM regression likely report *P*-values that are too low, and we believe CWM should be avoided entirely. If CWM is used, permutation tests should be performed to check type I error rates. MLM2 should be used instead, ideally with bootstraps to check *P*-values.

We have not addressed the issue of phylogenetic non-independence among species, and this can be an additional source of type I errors. Li and Ives (2017) show that if there is phylogenetic signal in the distribution of species among sites, such that closely related species are more likely to reach high abundance in the same sites, then type I error rates can be falsely inflated. Type I error control is particularly poor when the trait in the analysis also shows phylogenetic signal among species. While Li and Ives (2017) show this effect of phylogenetic non-independence using MLM2, we suspect that it also arises in MLM1 and CWM regressions for the same reasons.

Multi-level models not only produce better statistical tests than CWM regressions, they are more flexible. An example is MLM2 that can be modified to include phylogenies (Li & Ives 2017), which is not an option for CWM regressions. Multi-level models also provide significantly more information than CWM regressions about the nuances of trait-by-environment interactions in a community dataset. For example, a CWM analysis could show that average plant height increases in response to soil fertility. However, MLM2 will show that this pattern is driven by a negative relationship between plant height and species occurrence on low fertility soils while there is no relationship between species occurrence and height on high-fertility soils (see Jamil *et al*. 2013 for more examples and sample code). Finally, multi-level models can be used to address questions such as whether phylogenetically related species are less likely to occur in the same community as a possible consequence of competitive exclusion (Ives and Helmus 2011), or whether traits of flowers or pollinators are more important for explaining patterns of plant-pollinator associations (Rafferty & Ives 2013, Hadfield *et al*. 2014). Analyses of trait-by-environmental interactions is just one of numerous examples illustrating benefits of multi-level modeling approaches (Bolker *et al*. 2009, Warton *et al*. 2015).

## Data accessibility

- Community data: archived in DRYAD (doi: xx.xxx/dryad.xxxx)
- R scripts: included in online supporting information

